# Understanding the Evolutionary Games in NSCLC Microenvironment

**DOI:** 10.1101/2020.11.30.404350

**Authors:** Ranjini Bhattacharya, Robert Vander Velde, Viktoriya Marusyk, Bina Desai, Artem Kaznatcheev, Andriy Marusyk, David Basanta

**Affiliations:** Department of Life Sciences, Shiv Nadar University, Greater Noida, Uttar Pradesh, India; Department of Cancer Physiology, H Lee Moffitt Cancer Centre and Research Institute, Tampa, FL, USA; Department of Molecular Medicine, University of South Florida, Tampa, FL, USA; Department of Biology, University of Pennsylvania, Philadelphia, PA, USA; Department of Computer Science, University of Oxford, Oxford, UK; Department of Integrated Mathematical Oncology, H Lee Moffitt Cancer Centre and Research Institute, Tampa, FL, USA

## Abstract

While initially highly successful, targeted therapies eventually fail as populations of tumor cells evolve mechanisms of resistance, leading to resumption of tumor growth. Historically, cell-intrinsic mutational changes have been the major focus of experimental and clinical studies to decipher origins of therapy resistance. While the importance of these mutational changes is undeniable, a growing body of evidence suggests that non-cell autonomous interactions between sub-populations of tumor cells, as well as with non-tumor cells within tumor microenvironment, might have a profound impact on both short term sensitivity of cancer cells to therapies, as well as on the evolutionary dynamics of emergent resistance. In contrast to well established tools to interrogate the functional impact of cell-intrinsic mutational changes, methodologies to understand non-cell autonomous interactions are largely lacking.

Evolutionary Game Theory (EGT) is one of the main frameworks to understand the dynamics that drive frequency changes in interacting competing populations with different phenotypic strategies. However, despite a few notable exceptions, the use of EGT to understand evolutionary dynamics in the context of evolving tumors has been largely confined to theoretical studies. In order to apply EGT towards advancing our understanding of evolving tumor populations, we decided to focus on the context of the emergence of resistance to targeted therapies, directed against EML4-ALK fusion gene in lung cancers, as clinical responses to ALK inhibitors represent a poster child of limitations, posed by evolving resistance. To this end, we have examined competitive dynamics between differentially labelled therapy-naïve tumor cells, cells with cell-intrinsic resistance mechanisms, and cells with cell-extrinsic resistance, mediated by paracrine action of hepatocyte growth factor (HGF), within *in vitro* game assays in the presence or absence of front-line ALK inhibitor alectinib. We found that producers of HGF were the fittest in every pairwise game, while also supporting the proliferation of therapy-naïve cells. Both selective advantage of these producer cells and their impact on total population growth was a linearly increasing function of the initial frequency of producers until eventually reaching a plateau. Resistant cells did not significantly interact with the other two phenotypes. These results provide insights on reconciling selection driven emergence of subpopulations with cell non-cell autonomous resistance mechanisms, with lack of evidence of clonal dominance of these subpopulations. Further, our studies elucidate mechanisms for co-existence of multiple resistance strategies within evolving tumors. This manuscript serves as a technical report and will be followed up with a research paper in a different journal.

## Introduction

In 1859, Darwin’s “The Origin of Species” revolutionized scientific thought. He identified natural selection for adaptive heritable traits as the mechanism responsible for the evolution of natural populations [5]. Principles of Darwinian evolution can be applied towards the understanding of changes observed during somatic evolution, which underlies initiation and progression of cancers. [9][19][24]. Selection-driven clonal expansion in competitive contexts is responsible for the outgrowth of cells bearing genetically and epigenetically mediated alterations in cell-intrinsic characteristics (such as the acquisition of partial independence from growth factors, unlimited replicative potential and loss of sensitivity to apoptotic signals), as well as phenotypic traits responsible for adaptation to evolving tumor microenvironment (such as avoidance of immune mediated elimination, ability to grow at reduced pH and lower nutrients/oxygen concentrations etc) [16][27].

Similarly, Darwinian principles apply to evolutionary changes that occur in tumor cell populations during therapies. Despite a continuous expansion of oncologist’s arsenal of drugs, capable of suppressing and reversing tumor growth, advanced cancers inevitable develop resistance, with the rare exception of some immune therapy cases. With advances in therapeutic options, drug resistance becomes the leading causes of cancer-related fatalities [1][7][17][18]. Administration of drugs/ chemotherapy creates new selective pressures that act both immediately on proliferation/survival of tumor cells, as well as by altering tumor microenvironment [6] Historically, most of the research efforts has been focused on understanding cell-intrinsic mutational and epigenetic mechanisms that mediate reduced sensitivity cells to therapies. However, reduced therapy sensitivity can also reflect the impact of paracrine factors that can be produced both by tumor cells and non-tumor cells within the tumor microenvironment.

Lung cancer is one of the leading causes of cancer-related fatalities in the world [2]. Non-small cell lung cancer (NSCLC) accounts for 85% of all lung cancer diagnoses [22]. A specific subtype of NSCLC is EML4-ALK positive NSCLC that possesses a characteristic fusion between the two genes-echinoderm microtubule-associated protein-like 4 (EML4) and anaplastic lymphoma kinase (ALK) gene [38][39]. This fusion enables the production of EML4-ALK protein that is responsible for unchecked growth and malignancy. This subtype account for approximately 5 % NSCLC patients [10][23]. ALK tyrosine kinase inhibitor alectinib is currently used as a first line of therapy in EML4-ALK positive NSCLC patients. Studies have shown that cancer cells can bypass growth inhibition from alectinib by activating the alternative MET pathway triggered by binding with hepatocyte growth factor (HGF) [26]. Where, in some cases, cancer cells can acquire the ability to express HGF, HGF is generally expressed at high levels by cancer associated fibroblasts within tumor microenvironment (fig 1) [21].

**Fig 1:**
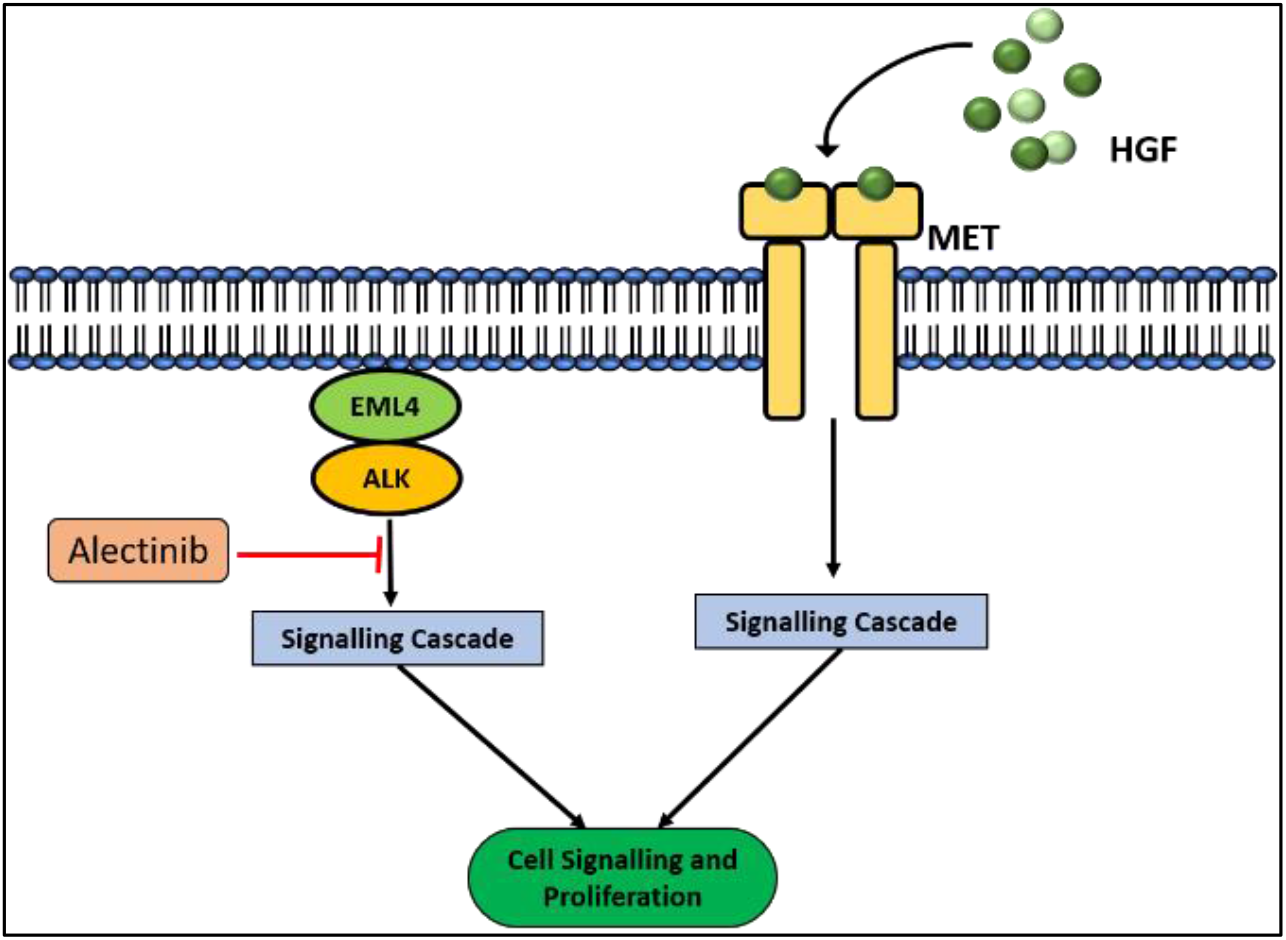
Inhibitory role of Alectinib on EML4-ALK positive NSCLC and activation of alternative c-MET pathway mediated by hepatocyte growth factor. The c-MET pathway activation mediates resistance to alectinib.

Relapse of tumor growth involves the selective expansion of therapy resistant subpopulations, which might either pre-exist or arise during the process of treatment. A growing body of evidence indicates that clinical resistance within the same tumors can be mediated by expansion of subpopulations with distinct resistance mechanisms. Expansion of interacting subpopulations with distinct phenotypic strategies can be viewed through the prism of evolutionary game theory, where the superior strategies win and obtain a reward (expansion) while inferior strategies pay a cost [12][20]. EGT has emerged as a branch of mathematics that enabled the extension of the concept of games (common in economic setups) to population biology [15]. EGT helps us account for the heterogeneity in the cancer microenvironment and their complex interactions [8][28][29]. In order to understand these dynamics, we set up simple *in vitro* assays using derivates of H3122 cell line, which has been a workhorse of experimental studies in ALK+ NSCLC [13]. In these assays, we examined competitive dynamics between therapy naïve cells, cells with acquired cell-intrinsic resistance to alectinib, and cells with cell extrinsic resistance, mediated by engineered ability to produce HGF.

We show that producers have the highest fitness in both drug exposed and unexposed conditions, in all heterotypic mixtures. They also provide a selective advantage to sensitive cells by activating the alternative MET pathway in them, thus mediating acquired drug resistance. This advantage is a linearly increasing function of the initial producer frequency up to a frequency of 0.4 after which the advantage reaches saturation. Our results also indicate that although the fitness of resistant cells is lower than that of sensitive cells in monoculture, they are fitter than sensitive cells in coculture. However, they do not actively participate in cross-talk with producer or sensitive cells. Our results elucidate the dynamics between three cellular strategies in a tumor microenvironment and how they mediate the emergence of resistance to a drug.

## Experimental Methods

Sensitive NSCLC cells (parental H3122), HGF overexpressing cells (H3122 cells engineered to produce HGF) and cells with evolved resistance to Alectinib (erAlec) were used to set up three sets of game assays. HGF overexpressing cell line was generated by transducing H3122 cells with pLenti6.3 lentiviral expression vector carrying isoform “#” of the human HGF gene. GFP and mCherry expressing cell lines were also generated via transduction using lentiviral constructs carrying the respective gene.Nuclear mCherry or GFP tagged cells (of each type) were allowed to grow until they reached 70-80% confluence. 2000 cells in total, were seeded in each well of a 384 well plate.

The three assays that were set up were (fig 3A)-

1. H3122 (mCherry) with H3122 o/e HGF (GFP),
2. H3122 (mCherry) with erAlec (GFP), and
3. H3122 (GFP) with erAlec (mCherry).

The initial seeding densities for the mCherry tagged cell line, for each assay varied from 0%, 2.5%, 5%, 10%, 15%, …., 95%, 100%, where the extremities (0 and 100%) represent monocultures, and the intermediate values represent coculture. For instance, 0% refers to 0% mCherry tagged cells and 100% GFP tagged cells. A seeding density of 10% would correspond to 200 mCherry tagged cells and 1800 GFP tagged cells. The total will always add up to 2000. Therefore, there are 21 initial seeding densities in total.

12 wells were seeded with one initial condition. After 24 hours, the cells were treated with DMSO (control) or different concentrations of Alectinib (0.125 uM, 0.25 uM, 0.5 uM). The 12 wells for each seeding density were now divided into 4 groups-3 each for the different drug concentrations and 3 for DMSO.

The plates were then placed in the Incucyte for 6 days. Time-lapse microscopy images were taken at intervals of 6 hours (fig 3B). The percent fluorescence confluence was analyzed on day 6. Exponential growth occurred until day 3 after which the populations began to saturate. The growth rates in the exponential phase were used as a measure of fitness.

## Game Assay

The replicator dynamic equations is a simple model for frequency-based evolution [25]. According to this model, the fitness of a strategy in a give population depends on the fitness of other strategies in the population composition and their prevalence. It measures the evolutionary success as the difference of fitness of a particular strategy and average fitness of the population [11][25].

For a population profile *X* where *x*_1_,.,.*x_n_* represent frequencies of strategies *1-n* in a given population

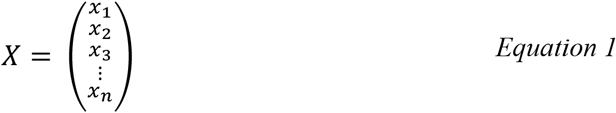

The per capita growth rate is given by :

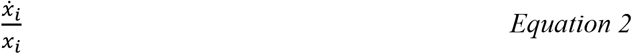

The average fitness of the population

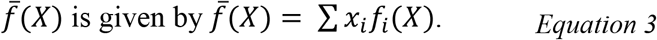

The replicator equation is given as-

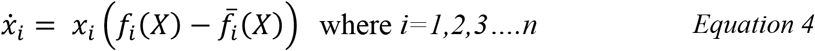

For a pairwise linear game, this can be represented as-

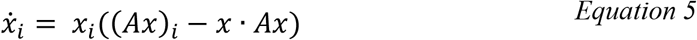

Where A is the pay-off matrix.

For

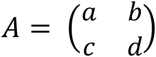

The replicator equations thus look like [13]-

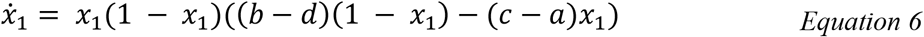

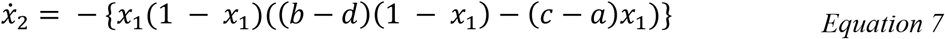

In our case, the population is comprised of two cell types from the three chosen cell lines: Producer (H3122 overexpressing HGF), resistant (erAlec), and sensitive (H3122). The output from the Incucyte was used to find the initial frequencies of each cell type using the formula-

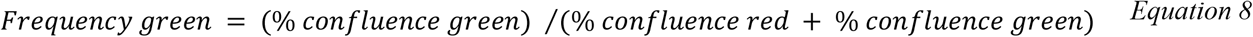

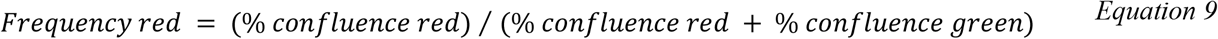

We assume that growth rate is a good estimation of cellular fitness. The individual fitness of each type is obtained from the growth rate in the exponential phase using a logarithmic regression (weighted by the inverse of error 1 on the growth rates) using the Theil Slope Estimator [13].

The growth rates were plotted against the initial experimental frequencies to characterize the trend. A linear trend appears to be a good estimate. The protective effect of HGF is more pronounced at lower frequencies and saturates above the frequency of 0.4. Therefore, we perform two regressions-one from 0-0.4 and one from 0.4-1 (initial frequencies). Although we perform two linear fits in two different ranges, we essentially obtain a single game with a non-linear fitness function. The fits were weighed against the inverse of error in growth rates (y-axis). The coefficients of these linear fits enable us to procure the payoffs for the payoff matrix for each assay. The coefficients of these linear fits enable us to procure the pay-offs for the pay-off matrix for each assay along with the errors on the entries in the same way as Kaznatcheev et al [13].

## Results

### The Games

#### Monoculture Behaviour

We found that the growth rate of sensitive cells is around 0.02 percent confluence/ hour. This growth rate dips down to around 0.008 percent confluence/ hour upon treatment with alectinib. Producers have a growth rate of 0.025 percent confluence/hour in both environments. We can see that producers have the highest growth rate and thus fitness in both conditions. We call the difference in the natural growth rates of sensitive and producer cells as the ‘benefit’ of production. The growth rate of resistant cells is approximately 0.015 percent confluence/hour. We call the reduction in its growth rate the ‘cost’ of resistance. This cost of resistance can be interpreted as an additional function (like energy allocation) that the cell needs to invest in to keep growing even in the presence of alectinib[14]. This means that they are lower in fitness than both producers and sensitive cells in untreated conditions but in the presence of alectinib, they are fitter than sensitive cells.

###### Producer VS Sensitive

- In both frequency ranges in control, the pay-off of producers interacting with producers is the highest. Producers derive more benefit from interacting with producers when compared to the sensitive population. Sensitive cells interacting with sensitive cells have the lowest pay-off.
- In alectinib, sensitive cells have the highest pay-off in the producer frequency range 0-0.4. This shows that at lower levels of HGF, the benefit to be derived is quite high.
- Note that in the frequency range 0.4-1, all strategies have a pay-off of greater than 2.0.

###### Resistant VS Producer

- The fit lines of resistant and producer are linear and parallel.
- Note that the pay-off of resistant population interacting with producer population is higher than that of the resistant vs resistant population.

###### Resistant VS Sensitive

- In both control and drug exposed condition, sensitive cells interacting with resistant cells have the lowest payoff.
- Although the fitness of resistant cells is higher than that of sensitive cells for most of the frequency range in control, sensitive cells appear to show a spike in fitness towards the end.

### HGF Enhances Fitness of Sensitive Cells

Heterotypic mixtures of two cellular phenotypes at a time, help us unravel how different strategies interact with each other [13][29].

We anticipated that the production of HGF by producer cells will benefit cells with other phenotypes [30][31] The presence of producers did not benefit sensitive cells in the absence of alectinib (fig 4.1, B) However, in the presence of alectinib, producers do benefit sensitive cells (fig 4.1, D) We found that sensitive cell growth rate increased with the proportion of producer cells. Once its growth rate reached its natural value (i.e. 0.02 confluence/hour) the fitness saturates. Thus, we see that the protection extended to sensitive cells by producers is linearly dependent on the frequency of producers up to a frequency of 0.4 after which it plateaus. The fitness of sensitive cells is much more sensitive to the amount of HGF present in the culture at lower frequencies of producers (0.005-0.2).

**Fig 2:**
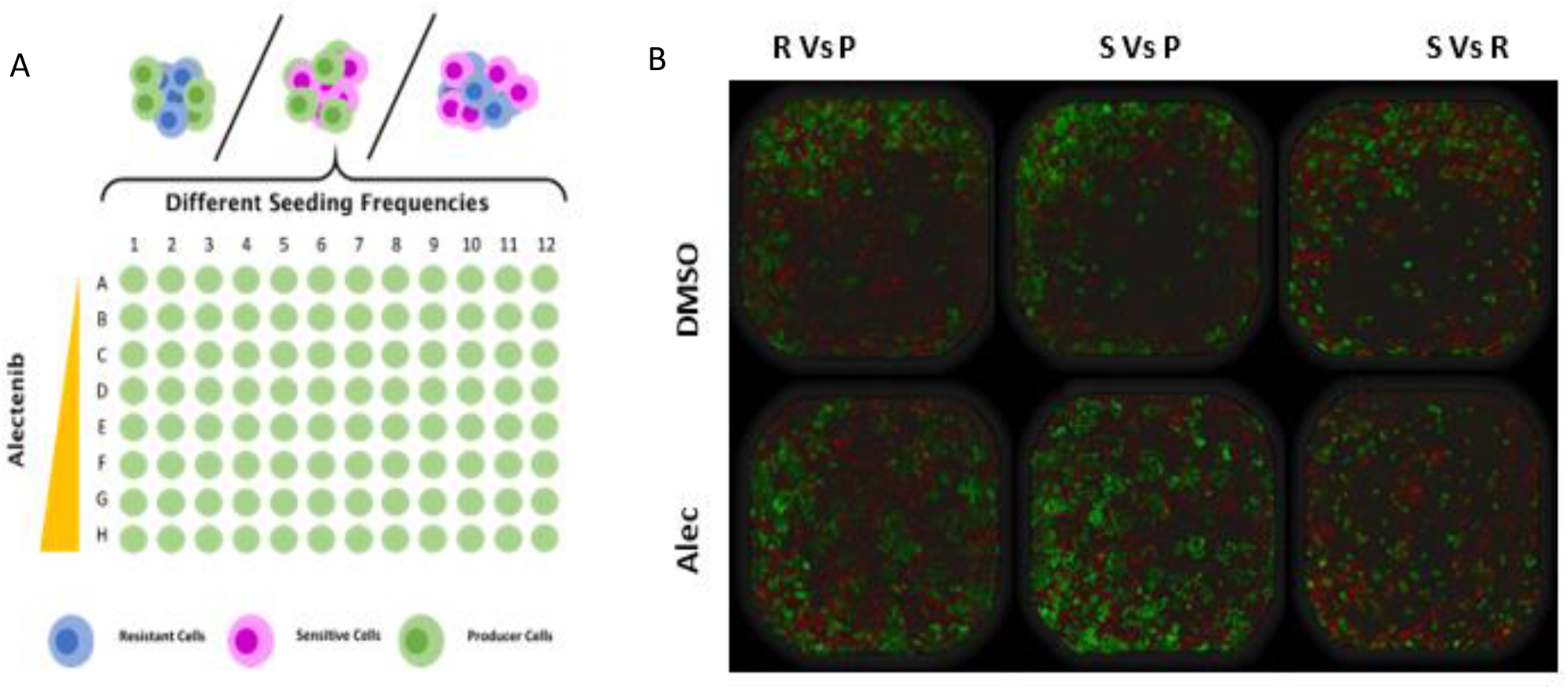
A: *In vitro* game assay strategy for setting up the three games. B: Images taken by the Incucyte in DMSO (control) and 0.5 μM alectinib after 4 h of seeding at 1:1 ratio

**Fig 4:**
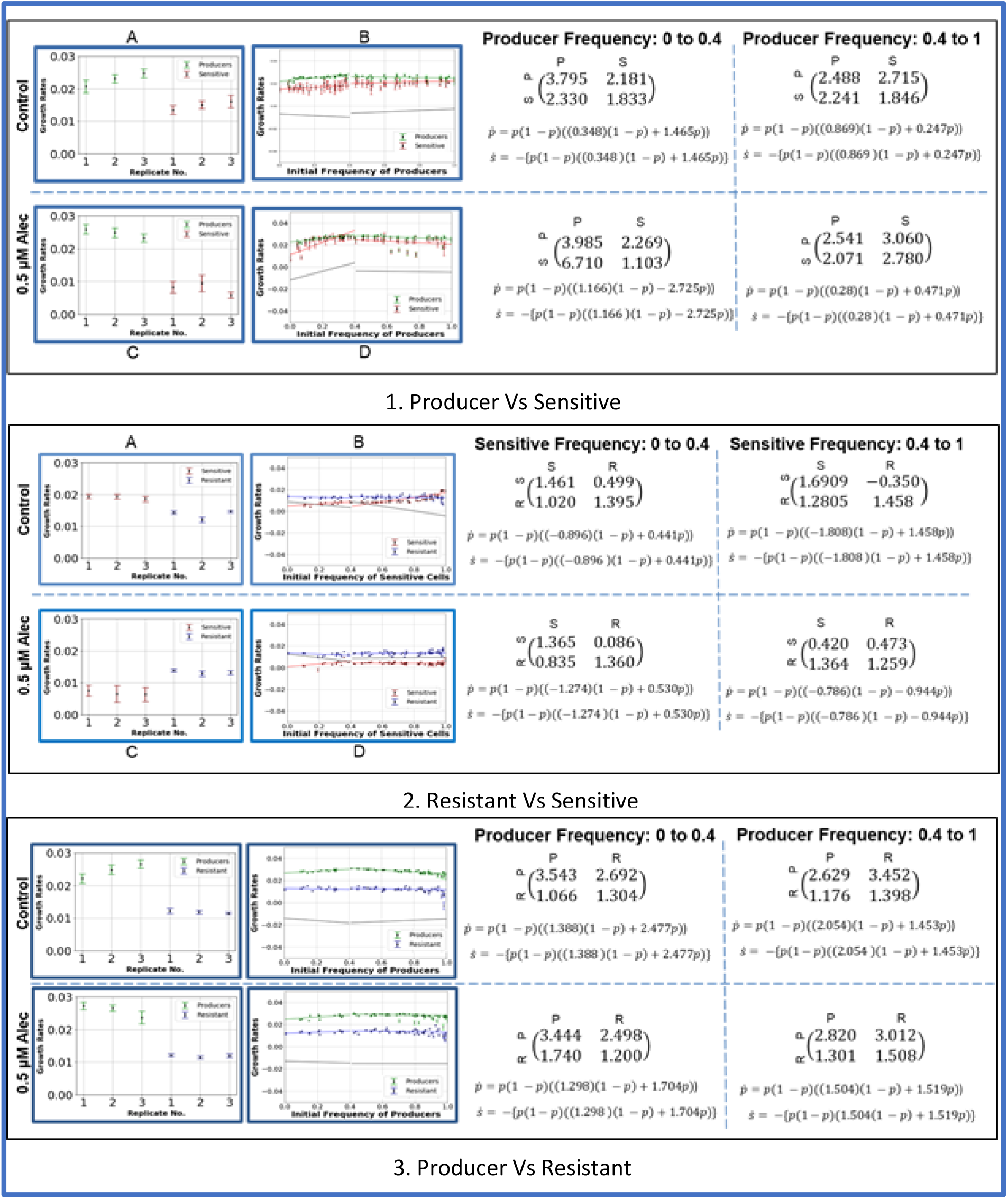
Summary of the three games in control and 0.5 uM Alectinib. Graphs A and C show the monoculture growth rates of the two cell populations in culture. Graphs B and D show the linear fits obtained between growth rate and frequency. Predicted gain in fitness in shifting from-1) producer to sensitive 2) resistant to sensitive 3)producer to resistant. Each entry *a_i,j_* in the matrix represents the payoff of strategy in row *i* interacting with strategy in column *j* where *i,J=1,2* Refer to fig S1, S2, and S3 for the entire summary.

However, we find that resistant cells do not derive any benefit from being in culture with producers. Their fitness neither increases nor decreases in the presence of producers. This could be because the intrinsic resistance mechanism is independent of the c-MET pathway and it continues to function regardless of the presence or absence of HGF. Alternatively, the cells could be physically incapable of growing any faster, regardless of the presence of growth factors.

### Resistance

The two games involving resistant cells-resistant vs producer and resistant vs sensitive demonstrate that resistant cells do not interact significantly with the other two cellular phenotypes. Their fitness remains constant irrespective of the coculture. This is in contrast with our previous results in Kaznatcheev et al [13] where we saw a clear dependence of fitness of resistant cells on the proportion of sensitive cells in coculture. Two key reasons that could be the cause of this difference are-1) the use of a different cell line and 2) use of a 384 well plate that reduces the space available for proliferation. Interestingly, in our experiments, we find that although the fitness of sensitive cells is lower than that of resistant cells for proportions below 0.9, its fitness jumps once the proportion of sensitive cells in culture is greater than 0.9, in control. In coculture with producer cells, producer cells are always fitter than resistant cells.

## Discussion

Paracrine acting factors like HGF can provide strong resistance to alectinib and other targeted therapies [21][30][31]. Whereas in accordance to the principle of evolutionary trade-offs it is commonly assumed that resistance is associated with a base-line fitness cost, we did not observe evidence of this trade-offs in our assays, at least in vitro. In contrast, cells with evolved intrinsic resistance to alectinib displayed a proliferative fitness penalty but were not affected by the presence of alectinib.

Studying these three games using a game-theoretic framework highlights the possibility that the payoffs received by individual cellular strategies can potentially be modified. For instance, we can experimentally modify producer cells to express higher or lower HGF which would alter the payoff matrices. Blocking the action of HGF receptor would help in enhancing the sensitivity of cancer to tyrosine kinase inhibitors. Therefore combination therapy with metformin might be useful [4]. A drug that inhibits the production of HGF would also make cancer more sensitive to treatment. Note that in this case, resistant cells will become fitter and thus the need to inhibit the resistant phenotype will also arise.

Our work underscores how paracrine effectors can modulate the fitness landscape of a tumor (fig 6). We illustrate this in *in vitro* system. The interactions will be much more complicated *in vivo.*

**Fig 5:**
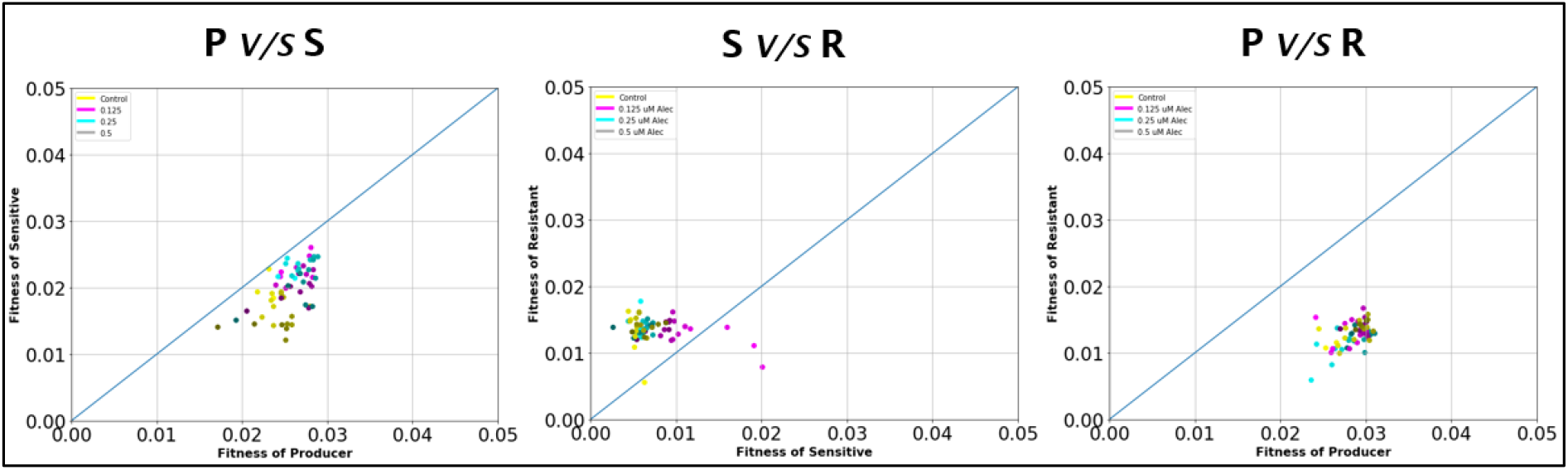
Comparative fitness of the different cellular populations in the pairwise games.

**Fig 6:**
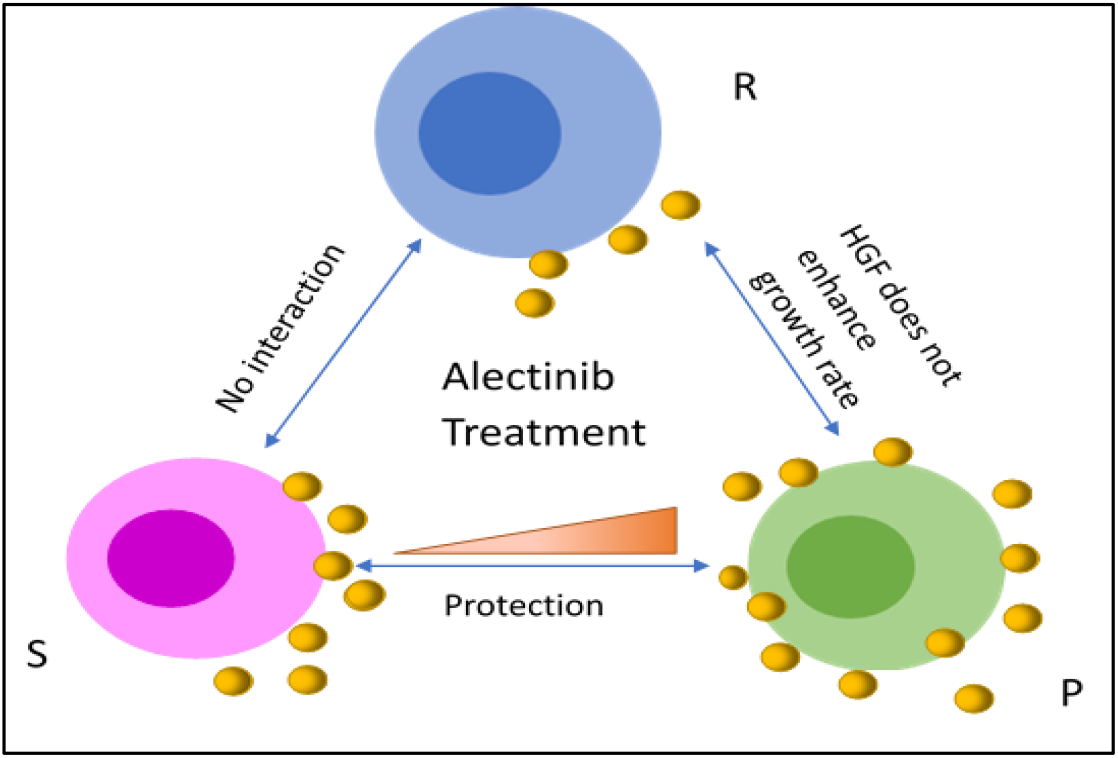
Dynamics between producers (P, green), resistant (R, blue), and sensitive (S, pink). The yellow circles represent HGF produced by P.

The ultimate goal of understanding dynamic relations in the tumor microenvironment using EGT is to treat the game instead of targeting individual players so that the entire evolutionary landscape can be tweaked for the benefit of the patient [13]. This manuscript serves as a technical report and will be followed up with a research paper in a different journal. We hope that this framework can be used to interpret complex interactions in tumors.

## Contributions

RB performed the experiments and analysis. RV did image analysis and helped design the assays. VM performed the experiments. BD helped with the experiments. AK guided with analysis of the data obtained and provided valuable feedback. AM and DB visualized the project, provided feedback, and helped with troubleshooting. RB wrote the manuscript. RV, AK, AM, and DB reviewed and provided feedback on the manuscript.

## Acknowledgements

RB acknowledges the Khorana Scholars Program for the fellowship to undertake this project. DB was partially supported by an NCI IMAG-MSM grant (U01CA202958). DB was also funded by PSON U01 CA244101. AM and DB were partially supported by the State of Florida award 20B06 (30-20450-99-01).

All the authors thank We would also like to thank Aakash Grover, Jack Edwards, Siddharth Choudhary, and Mark Laurie for their insightful feedback and discussions.

