## Supplementary Figures for "Understanding the Evolutionary Games in NSCLC Microenvironment"

**Summary of the Game**

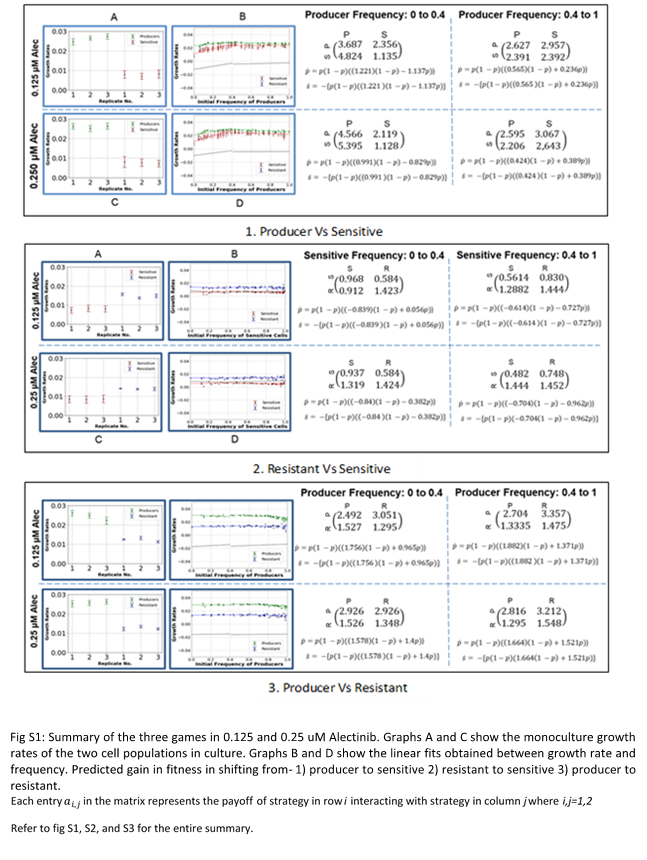

**Frequency Graphs**

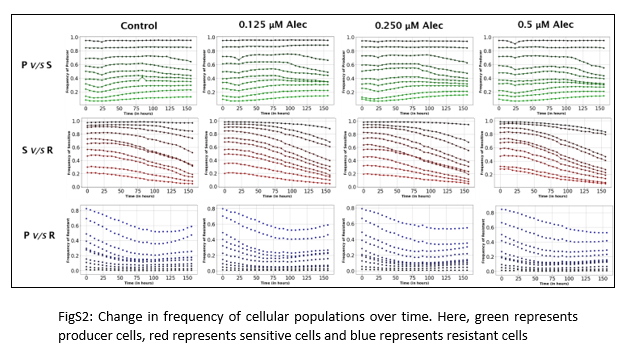

**Confluence Graphs**

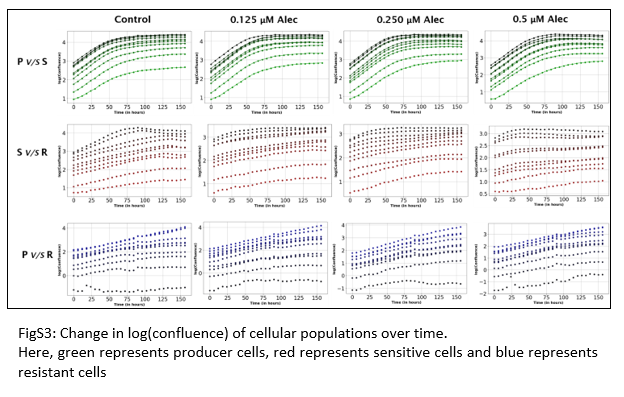

**Residual Plots**

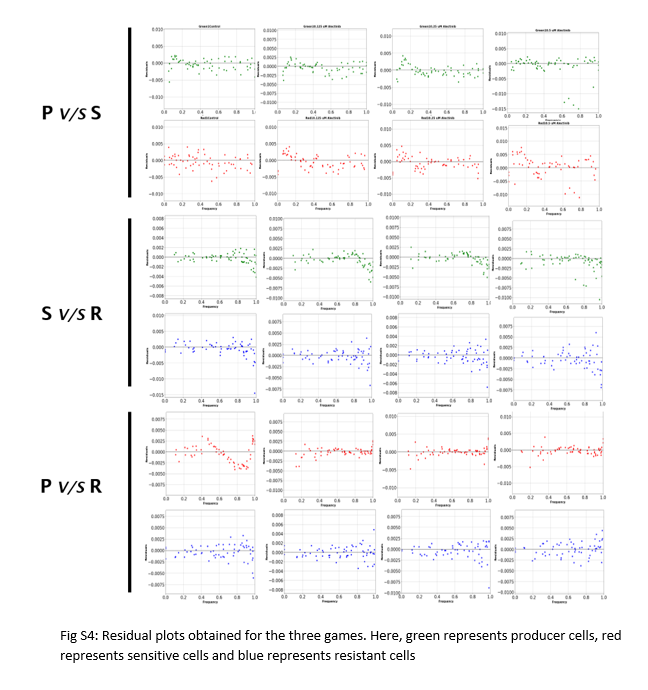
